# Identifying Lymph Node Metastasis-related Factors in Breast Cancer using Differential Modular and Mutational Structural Analysis

**DOI:** 10.1101/2022.09.06.506724

**Authors:** Xingyi Liu, Bin Yang, Xinpeng Huang, Wenying Yan, Guang Hu

**Affiliations:** Center for Systems Biology, Department of Bioinformatics, School of Biology and Basic Medical Sciences, Soochow University, Suzhou 215123, China

## Abstract

Complex diseases are generally caused by disorders of biological networks and/or mutations in multiple genes. Network theory provides useful tools to study the underlying laws governing complex diseases. Within this framework, comparisons of network topologies, including the node, edge, and community, between different disease states can highlight key factors within these dynamic processes. Here, we propose a differential modular analysis approach that integrates protein-protein interactions with gene expression profiles for modular analysis, and introduces inter-modular edges and date hubs to identify the “core network module” that quantifies the significant phenotypic variation. Then, based on this core network module, key factors including functional protein-protein interactions, pathways, and drive mutations are predicted by the topological-functional connection score and structural modeling. We applied the approach to analyze the lymph node metastasis (LNM) process in breast cancer. The functional enrichment analysis showed that both inter-modular edges and date hubs play important roles in cancer metastasis and invasion, and in metastasis hallmarks. The structural mutation analysis suggested that the LNM of breast cancer may be the outcome of the dysfunction of rearranged during transfection (RET) proto-oncogene-related interactions and the non-canonical calcium signaling pathway via an allosteric mutation of RET. We believe that the proposed method can provide new insights into disease progression such as cancer metastasis.

**Author summary:** Metastasis is the hallmark of cancer that is responsible for the greatest number of cancer-related deaths. However, it remains poorly understood. PPI networks not only provide a static picture of cellular function and biological processes, but also have emerged as new paradigms in the study of the dynamic process of disease progression, including cancer metastasis. Herein, a network-based strategy was proposed based on the integration of expression profiles with protein interactions, by filtering with “date hubs” and “inter-modular edges”, demonstrating that different network modules may provide robust predictors to represent the dynamic mechanisms involved in metastasis formation. Furthermore, the mapping of protein structure and mutation data on the network module level provides insight into signaling mechanisms; helps understand the mechanism of disease-related mutations; and helps in drug discovery. The application of our method to study the LNM in breast cancer highlights network modules defining protein communities that respond to therapeutics, and the implications of detailed structural and mechanistic insight into oncogenic activation and how it can advance allosteric precision oncology.

## Introduction

Complex diseases, especially cancers, occur under the combined effects of many factors, and are generally considered to be caused by the disorder of molecular networks or biological systems.[1] Metastasis is the hallmark of cancer and it is responsible for the greatest number of cancer-related deaths,[2, 3] constituting the primary cause of death for >90% of patients with cancer. The activation of cancer metastasis has three distinguishing features: location-dependence, environmental interaction, and a dynamic selection process. Regional lymph nodes (LNs) are often the first sites of breast cancer metastasis and the first to encounter the host immune surveillance mechanisms intended to destroy foreign invaders. Lymph node metastases (LNMs) in cancer patients are associated with tumor aggressiveness, poorer prognoses, and the recommendation for systemic therapy.[4] For example, LNM is one of the most important independent risk factors that can negatively affect the prognosis of breast cancer.[5] Due to the dynamic nature of LNMs,[6] systems biology-based approaches may provide useful tools for understanding their molecular mechanisms and guiding treatment.

Benefiting from the advances of network science and high-throughput biomedical technologies, studying biological systems from network biology has attracted much attention in recent years. Networks have long been central to our understanding of biological systems, in the form of linkage maps among genotypes, phenotypes, and the corresponding environmental factors.[7] With the tremendous increase in human protein interaction data, the protein-protein interaction network (PPIN) approach commonly used to understand the molecular mechanisms of disease, and in particular to analyze cancers.[8] However, PPINs merely provide a static snapshot of the molecular interactions within a tissue, whereas biological systems are highly dynamic. Thus, differential network analysis can be used to study the dynamic properties of networks related to cancer metastasis and to highlight network changes between conditions.[9] Differential network methods developed to date differ in the entities and measures that they compare.[10] Node-based methods focus on differences in node-related measures, such as node connectivity.[11] Interaction-based methods focus on differences in the context-specific weights associated with each interaction.[12] Furthermore, integrating co-expression data with differential network analysis can reveal the dynamic context of gene expression profiles.[13, 14] Such differential co-expression networks are useful tools to identify changes in response to an external perturbation, such as mutations predisposed to cancer progression, and to identify changes in the activity of gene expression regulators or signaling.

Three-dimensional protein structural data at the molecular level are pivotal for successful precision medicine. Such data are crucial not only for discovering drugs that act to block the active site of the target mutant protein but also for clarifying to the patient and the clinician how the mutations harbored by the patient work.[15] In oncological research, structure-based methods including driver prediction, computational mutagenesis, (off)-target prediction, binding site prediction, virtual screening, and allosteric modulation analysis, can be highly beneficial for addressing the diversity of cancer hallmarks.[16] Recently, a structure-function-based approach was shown to improve the prediction of drug sensitivity in Epidermal growth factor receptor (EGFR)-mutant non-small cell lung cancer.[17] Actually, linking structural information with PPI network or biological pathway information is useful for predicting the genotype-phenotype relationship, providing insight into signaling mechanisms, helping to understand the mechanism of disease-related mutations, and helping in drug discovery. Such an approach has been used for the detailed analysis of cancer and the cancer metastasis-related PPI binding interface,[18, 19] and to delineate the mechanism of oncogenic mutations and single nucleotide polymorphism mutations in inflammation and cancer.[20]

Worldwide, breast cancer is the leading cause of cancer-related deaths in women.[21] Despite the decline in breast cancer mortality and recent advances in targeted therapies and combinations for the treatment of metastatic diseases, metastatic breast cancer remains the second leading cause of cancer-related death among women in the United States.[22] The metastasis of breast cancer mainly occurs through the lymphatic system. Therefore, understanding the biological process and mechanism of LNM in breast cancer will help guide the treatment of breast cancer and improve the prognosis of patients.[23] Although several studies have focused on the identification of disease markers in metastatic breast cancer from the perspective of genome-wide expression profiles[24] and comparative analysis of PPI networks,[25] the molecular understanding of LNM in breast cancer is still very poor. In this paper, based on the idea that network modules serve as a more robust indicator of cancer prognosis,[26, 27] we propose a differential modular analysis approach to identify key network modules correlated with LNM in breast cancer. Firstly, weighted PPI networks for non-LNM and LNM were constructed by incorporating human interactome and gene expression data. Then, based on network modular analysis, “inter-modular edges” and “date hubs” were introduced to detect the altered modularity of PPI networks, which may correspond to the key dynamic region in LNM. Finally, we evaluated the importance and potential application of the core network module by mutational structural analysis at both the edge and node levels. We hope that this study provided a novel perspective for the analysis of mutation effects to facilitate network-guided precision medicine. An overview of our approach is shown in Fig 1, and the source code used in this paper can be found at https://github.com/CSB-SUDA/DMA.

**Fig 1.**
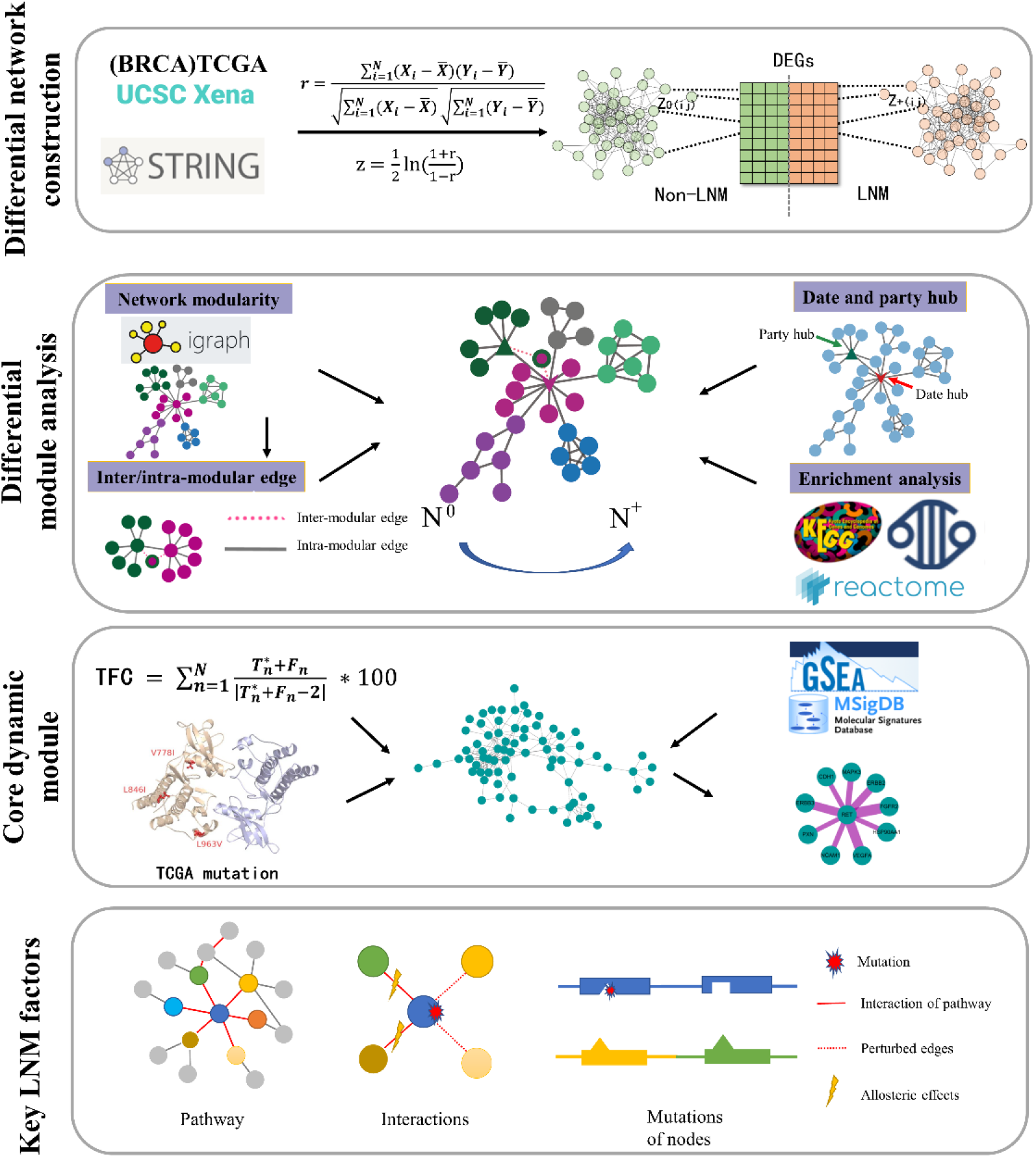
Overview of our approach. First, two weighted PPI networks for both LNM and non-LNM in breast cancer were constructed, whose topology is based on the STRING database and DEGs of LNM and non-LNM, weighted by the r-z transformation of co-expression data. Second, module analysis was performed for the two weighted PPI networks, and inter/intra-modular edges and date/party hubs, as well as KEGG, Reactome, and cancer hallmark enrichment analysis were introduced to compare the topology and biological functions of different modules. Third, the core network module for breast LNM was further characterized by assessing TFC scores for edges, structural modeling and mutation mapping, and GSEA enrichment analysis. Fourth, three key factors were predicted: allosteric mutations, key PPIs, and key pathways for LNM in breast cancer.

## Materials and Methods

### Data collection and pre-processing

RNA-seq data on breast cancer (BRCA) from The Cancer Genome Atlas (TCGA) with clinical information were retrieved from UCSC XENA (https://xena.ucsc.edu/). According to the extent of the lymph node metastases (LNM) of the clinical information, BRCA patients were divided into N^0^ (non-LNM) and N^+^ (LNM) groups and missing information was discarded. Complete classification information is provided in Table 1. The value of expression was converted to log_2_(TPM+1). We selected the genes with the highest median absolute deviation of 75% in the expression profile screening mean, at least larger than 0.01. Data analysis was performed to analyze the differences between the N^+^ and N^0^ groups using the package “limma” in R, and P-value < 0.01 was set as the cut-off to screen for differentially expressed genes (DEGs).

**Table 1.** Group of BRCA samples for regional lymph node metastasis.

| Extent of LNM | The number of samples | Group |
| --- | --- | --- |
| null | 2 | discarded |
| N0 | 333 | N<sup>0</sup> |
| N0(i-) | 154 | N<sup>0</sup> |
| N0(i+) | 28 | N<sup>0</sup> |
| N0(mol+) | 1 | N<sup>0</sup> |
| N1 | 126 | N<sup>+</sup> |
| N1a | 170 | N<sup>+</sup> |
| N1b | 33 | N<sup>+</sup> |
| N1c | 2 | N<sup>+</sup> |
| N1mi | 36 | N<sup>+</sup> |
| N2 | 56 | N<sup>+</sup> |
| N2a | 64 | N<sup>+</sup> |
| N3 | 26 | N<sup>+</sup> |
| N3a | 49 | N<sup>+</sup> |
| N3b | 3 | N<sup>+</sup> |
| N3c | 1 | N<sup>+</sup> |
| NX | 20 | discarded |

Footnote: TNM Classification of Malignant Tumors: N0, N0(i-), N0(i+), N0(mol+), N1, N1a, N1b, N1c, N1mi, N2, N2a, N3, N3a, N3b, N3c, and NX; N<sup>0</sup> denotes the non-LNM group, and N<sup>+</sup> denotes the LNM group. We discarded the means not participating in the group.

To verify the prognostic prediction performance of LNM, Kaplan-Meier survival analysis and the log-rank test were performed to identify the prognostic significance of LNM between the N^0^ and N^+^ samples. Kaplan-Meier survival curves and log-rank tests were executed using the R packages “survival” and “survminer”. The mutation data of the corresponding sample using the MuTect2 pipeline were downloaded to compare the tumors to a pool of normal samples to find somatic variations. Somatic mutations were analyzed with the R package “maftools” and visualized in a waterfall plot.[28]

### Differential interaction networks construction

The two differential weighted PPI networks under specific conditions (non-LNM and LNM) were built with the following steps. First, the protein-protein interaction network for *Homo sapiens* was retrieved from the STRING database,[29] filtering interactions by a combined score > 0.4. Second, DEGs between N^0^ and N^+^ were used to screen the interactions to form the topological structures of the two target networks. Finally, the Pearson’s correlation coefficients (PCCs) between the gene expressions were transformed by Fisher r-z transformation,[30] and its absolute value as the weights of the two networks, defined as:

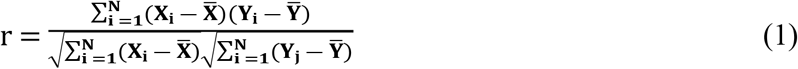

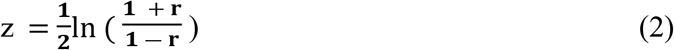

where X_i_ denotes the sample gene expressed value indexed *i*, and analogously for Y_i_, and 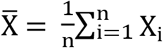(the gene expression mean value), and analogously for 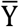. The absolute value of *z* is the weight of interaction between *i* and *j* genes.

### Differential modular analysis

#### Module detection

Integrated co-expression networks were then clustered using the multi-level modularity optimization algorithm.[31] The method is based on the modularity measure and a hierarchical approach, and the resolution parameter is set to 1. Here, only modules larger 10 nodes were considered. Module detection was performed with the R package “igraph” and represented using Cytoscape.[32] Next, we calculated the similarity between two modules using the Jaccard similarity coefficient, defined as

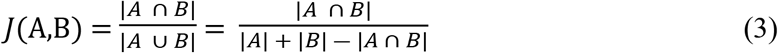

where A and B are the node sets of two different modules.

#### Edge knockout experiment

In the edge knockout experiment, two commonly topological measures were calculated:[33] betweenness, which measures the information flow through networks; and the characteristic path length (CPL), which is the average of the shortest path between all nodes in a network. The change of the two topological measures was applied to systematically access the robustness of the network by removing the equivalent number of different edges.

#### Date and party hubs

The average Pearson’s correlation coefficient (aPCC) was calculated for each interaction in the PPI network based on the co-expression of two interacting genes. Then, two types of hub genes were defined as date hubs and party hubs with low and high aPCC values, respectively.[26, 34] Random sampling of the PCC was used to ascertain that the observed edges were nonrandom.

#### Topological-functional connection

The topological-functional connection (TFC) was used for the prioritization of PPIs. The TFC is the integration of the edge betweenness and the gene ontology (GO) semantic similarity,[35] as a new edge measure:

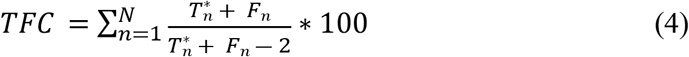

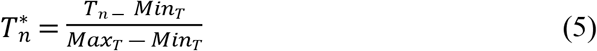

where *N* represents the number of interactions, and *T*_*n*_ and *F*_*n*_ represent the edge betweenness and GO semantic similarity, respectively, of interaction *n*. Thus, the TFC score is proposed to identify key protein interactions by integrating network topology and biological characteristics.

#### Enrichment analysis

Four types of functional enrichment analysis were used in our work. To identify the significant biological pathways of each module, we used the R package “clusterProfiler”[36] to perform Kyoto Encyclopedia of Genes and Genomes (KEGG) pathway enrichment.[37] Terms with an adjusted P-value < 0.01 were considered significant. Gene set enrichment analysis (GSEA) was performed to investigate the particular pathway across the whole expression profile, using the MSigDB database[38] and the R package “GSVA”.[39] Genes were sorted according to the logFC in the results of the N^0^ and N^+^ group difference analysis.

Unlike common Reactome enrichment analysis based on gene set, we performed another pathway enrichment based on interaction (or edge) annotations from the Reactome database.[40] The annotations are inferred between all protein components of a complex. To determine whether the gene set corresponded to cancer hallmarks, the hallmark annotations from the Catalogue of Somatic Mutations in Cancer (COSMIC) data resource were used.[41] Both of these enrichments were performed by using the R package “clusterProfiler.” Terms with a P-value < 0.05 were considered significant.

#### Structural modeling of protein-protein interactions

Protein structures were obtained from the Protein Data Bank (PDB). We used PRISM[42, 43] (Protein Interactions by Structural Matching) to predict the structures of protein-protein interactions (PPIs) and the effect of the mutations on interactions. Predicted protein complexes were ranked by FiberDock[44] according to their energies, and complexes with the lowest binding energy were selected to evaluate the effects of mutations on PPIs. Thus, the effect of a mutation on PPI is defined as:

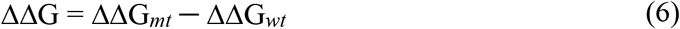

where ΔΔG_*mt*_ and ΔΔG_*wt*_ are the lowest binding energies of the mutant and wild-type complexes, respectively; and ΔΔG is the binding energy change caused by a single mutation.

#### Allosteric effects of mutations

The structure-based statistical mechanical model of allostery was used to obtain a direct estimate of the allosteric effects caused by the single mutation, by using AlloSigMA.[45] In the approach, two types of mutations were defined: UP-mutation, which models the situation of an actual mutation to a bulky residue with over-stabilizing effects on the local contact network; conversely, DOWN-mutation models the destabilization of the residue’s contact network similarly to Ala/Gly-like mutations. The allosteric free energy of the residue quantifies the strength and sign of allosteric communication associated with the mutation, where positive and negative signs correspond to a kinetic increase (local destabilization) and decrease (local stabilization), respectively.

## Results

### N^0^ and N^+^ PPI networks present different modular structures

According to the data in Table 1, 1579 differentially expressed genes (DEGs) were obtained, including 1,299 up-regulated and 280 down-regulated genes in N^+^ compared with N^0^. A volcano plot of the DEGs is shown in S1a Fig, in which *MISP, BLVRA, RNF223, KRT8, PTK6, PQLC3, KRT18, BCL2L14, ETV6*, and *LMO4* were the 10 most significantly differentially expressed genes. In addition, a Kaplan-Meier model for N^0^ and N^+^ was constructed, and we observed that the survival status of the lymph node metastasis group was significantly worse than that of the non-lymph node metastasis group, with a log rank test P-value < 0.0001 (S1b Fig). This result also confirmed that lymph node metastasis could be an independent predictor of survival in breast cancer.

Based on these DEGs, the corresponding topological topology of the LNM-related PPI network was generated, with 1,516 nodes and 8,286 edges. The absolute value of the Pearson coefficient between the expressions in the N^0^ and N^+^ groups was introduced as network weights, and then two different weighted PPI networks for LNM-related PPI networks were finally constructed, defined as the N^0^ and N^+^ PPI networks. As such, these two networks could preliminarily reflect the dynamic nature of the LNM process. Next, the modular detection algorithm was used to discover the sub-network structures and functions of the PPI networks. From a global perspective, the N^0^ and N^+^ PPI networks can be separated into 15 and 17 modules (Fig 2a and S1c Fig), respectively. Almost all modules tend to rewire, and smaller modules can be obtained in the N^+^ PPI network. A heat map of module similarity further quantifies this result; that is, the topological properties of only three modules are preserved (Jaccard similarity > 0.6), but the remaining modules have been changed (S1d Fig).

**Fig 2.**
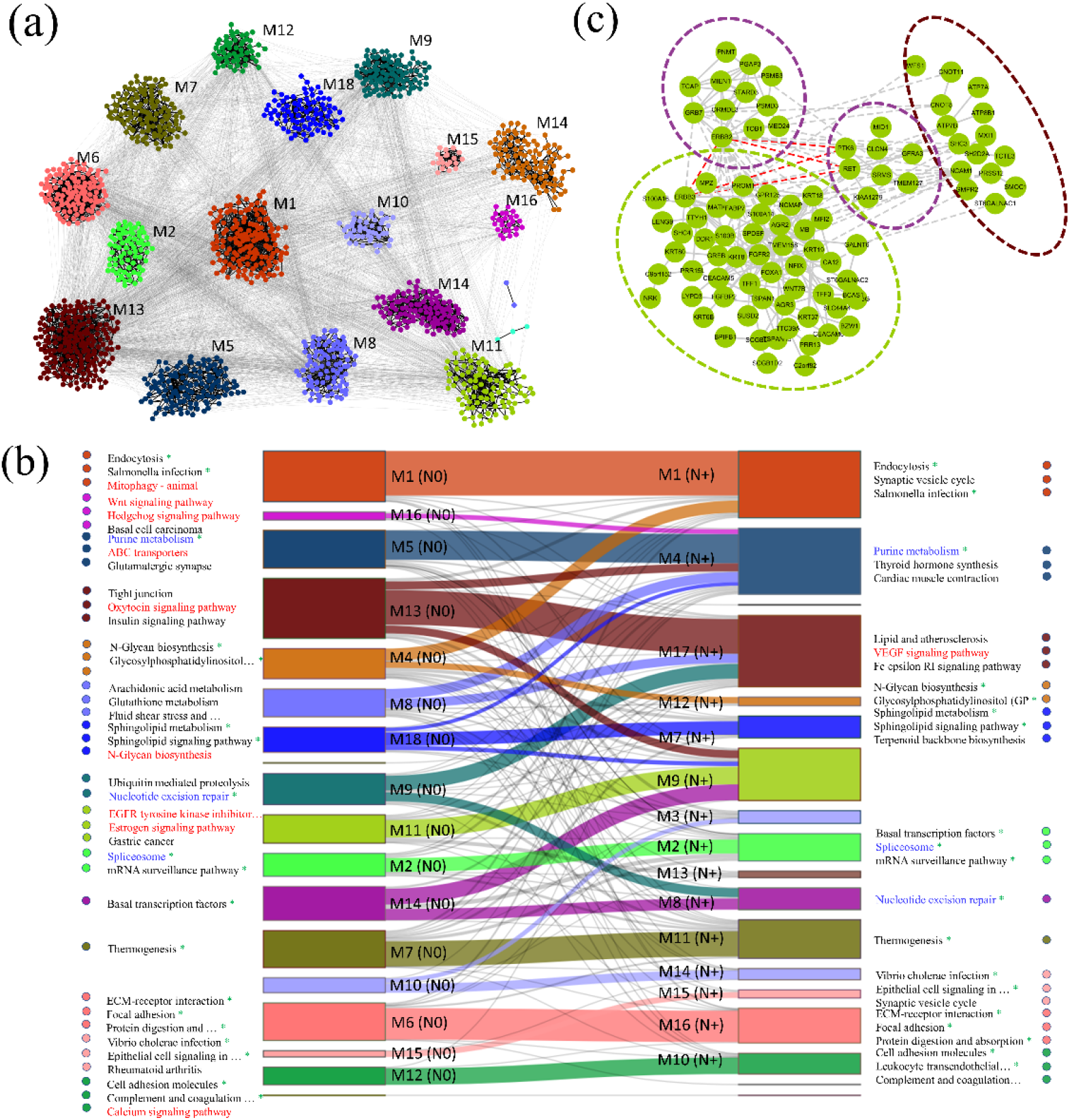
Overview of modular structures and functions of LNM-related PPI networks. (a) Modular structure of the N^0^ network, where nodes are colored according to different modules. (b) Assignment flow to KEGG pathways from modules in the N^0^ to N^+^ networks, shown in a Sankey diagram. Each module is annotated by the three most significant pathways with colored beads, or by all related pathways if the enriched pathways number fewer than three. Shared pathways by the N^0^ and N^+^ modular networks are marked by green asterisks, and pathways colored by red and blue indicate cancer metastasis-related biological pathways. (c) Example of ERBB2-related interactions in the modular structures between the N^0^ and N^+^ networks. Dashed red lines correspond to the observed interactions that locate between different modules of the N^+^ network and inside a module of the N^0^ network.

The Sankey diagram in Fig 2b shows the assignment flow from N^0^ modules to N^+^ modules. To further compare the biology underlying the modular change, we examined their related biological pathways. We found that most cancer-occurrence and progression-related basic pathways were preserved (green asterisks), including purine metabolism, nucleotide excision repair, and spliceosome, as highlighted in blue font. Furthermore, some statistically significant pathways in the N^0^ modular network were not significant in the N^+^ state (highlighted in red font). Among these pathways, we found that most corresponded to signaling transformation and cancer metastasis. In particular, the imbalance of the calcium signaling pathway in breast cancer will lead to the migration, invasion, proliferation, tumorigenicity, or metastasis of cancer cells. The expression of some key proteins in breast cancer is closely related to its progression. For example, oncogenic receptor tyrosine kinase *ERBB2* (also called *HER2*) is overexpressed in approximately 20% of breast cancers, resulting in ligand-independent dimerization and activation. The preliminary modular analysis shows that ERBB2-ERBB3, ERBB2-RET, and ERBB2-RTK6 interactions are involved in the same module of the N^0^ network, but between different modules of the N^+^ network (Fig 2d). Due to the importance of ERBB2 in breast cancer, the change in ERBB2-related interactions during LNM may be useful as an indicator of breast cancer metastasis.

The comparison of the network modules of N^0^ and N^+^ PPI networks demonstrated that most biological functions are conserved during LNM, although some module evaluation was also detected. Accordingly, further study of these different modules will help us understand the LNM mechanism and facilitate the prediction of key modules, interactions, and genes involved in cancer metastasis.

### Intermodular edges correspond to network signaling and cancer metastasis

To further characterize how modular structures change during LNM, we classified the edges or PPIs involved in dynamic changes into two types: 1) inter-modular edges, whose interacting nodes are located within the same module in the N^0^ PPI network, but within different modules in the N^+^ PPI network, or vice versa; and 2) intra-modular edges, in which the two interacting nodes are always located within the same module or between different modules. In all, 1,546 inter-modular edges and 6,740 intra-modular edges were identified in the LNM-related network dynamic process. Topological and functional analyses of these edges were then performed in terms of edge knockout experiments, as well as Reactome pathway enrichment analysis based on edge annotation.

In the edge knockout experiment, we calculated the average betweenness and average shortest path of each remaining network by systematically removing two types of edges randomly based on the N^0^ network, with a gradient of 20 and 200 repetitions. The average betweenness of the interaction network decreased more quickly by removing inter-rather than intra-modular edges (Fig 3a), while the average shortest path increased more quickly by removing inter-rather than intra-modular edges (Fig 3b). Betweenness measures the information flow through networks, with high betweenness indicating high biological signaling ability. The characteristic path length (CPL) indicates the network entropy, meaning that a biological network system with a higher CPL is more chaotic. Both topological measures showed that their values were more sensitive to intermodular edges, indicating that inter-modular edges contributed more to maintaining the global connectivity of the network, and played key roles in network signaling.

**Fig 3.**
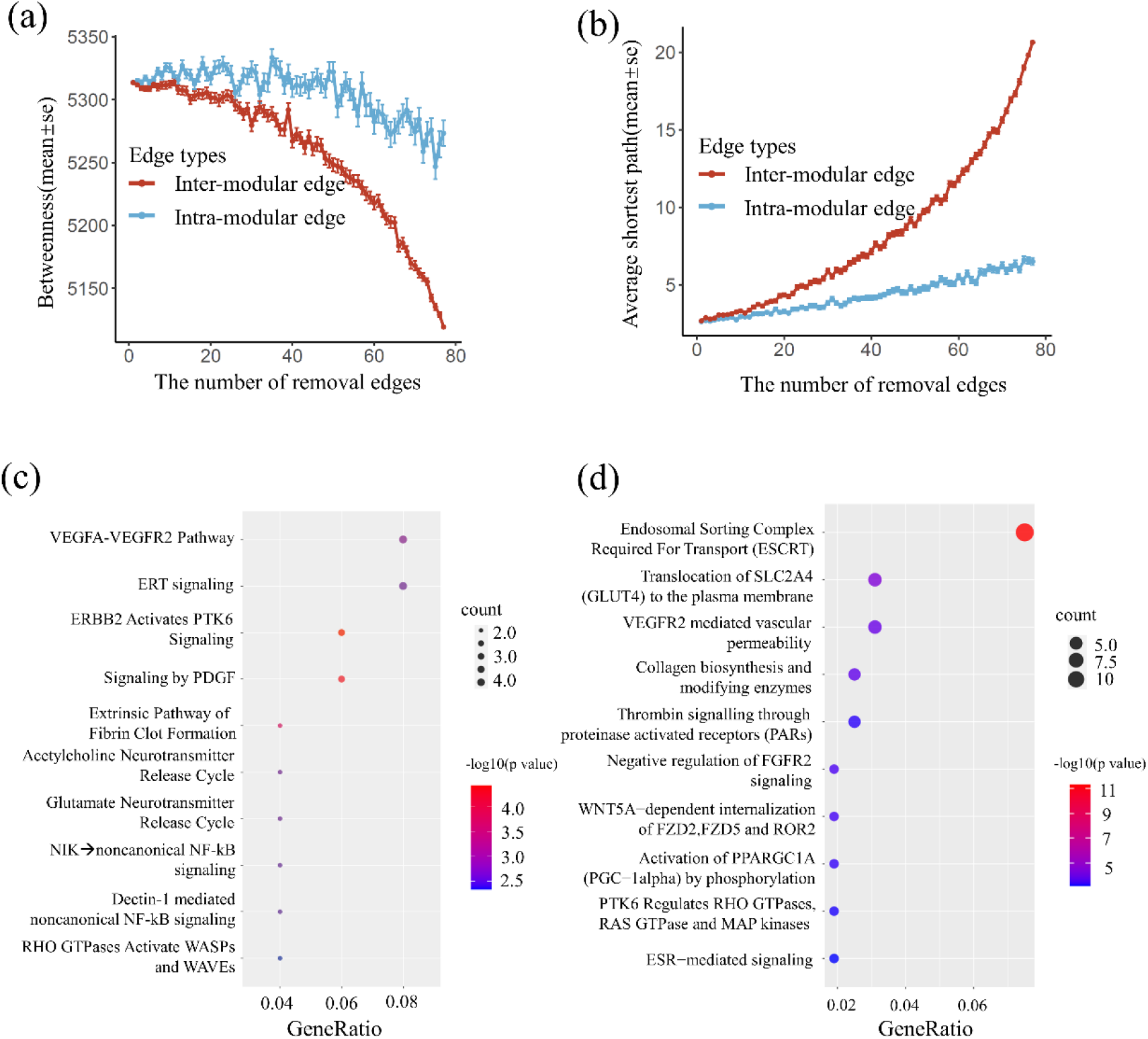
Topological and functional analysis of inter/intra-modular edges. (a) Network betweenness as a function of removing equivalent numbers of inter-modular and intra-modular edges. (b) The characteristic path length of the network as a function of removing equivalent numbers of inter-modular and intra-modular edges. Significantly enriched Reactome pathways of edges in the (c) inter-modular and (d) intra-modular groups

To identify specific biological functions, pathway enrichment of these two types of edges was performed based on “edge annotation” in the Reactome database. As shown in Fig 3c, inter-modular edges were significantly enriched with several cancer emergence- and development-related pathways, including the most significant VEGFA-VEGFR2 pathway, related to angiogenesis, followed by RET signaling, ERBB2 Activates PTK6 Signaling and Signaling by PDGF. On the other hand, significant Reactome pathways of the intra-modular edges are mainly involved in transport, metabolism, protein modification, growth and development, and signal communication, such as Endosomal Sorting Complex Required For Transport (ESCRT), Translocation of SLC2A4 (GLUT4) to the plasma membrane, VEGFR2-mediated vascular permeability, collagen biosynthesis, and modifying enzymes (Fig 3d). Together, the intermodular edges contributed to the major events implicated in the network signaling pathway and served as attractive targets for inhibiting LNM.[46]

### Date hubs reveal the invasion and metastasis hallmarks

Topological analysis of the PPI networks revealed that most of the proteins were connected to relatively few, highly connected proteins, termed hub proteins. Hubs in PPI networks have been classified into party and date hubs based on the co-expression of the interacting proteins.[2] Whereas date hubs display low co-expression with their partners, party hubs have high co-expression. It has been proposed that date hubs were global connectors, whereas party hubs were local coordinators. Date hubs tended to have transient interactions due to their low average co-expression correlation with their interaction partners, which not only played a role in connecting biological modules to each other but also show more dynamic properties.

Here, we defined nodes with a degree > 20 in PPI networks as hub nodes. The average Pearson’s correlation coefficients (PCCs) of the edges related to these hub nodes in the N^0^ network and the N^+^ network were calculated. Then, the date hubs and party hubs were defined as having lower PCCs and higher PCCs by the median of these values. To investigate the functions of these types of hubs, we first compared the relationship between hub proteins and inter/intra-modular edges. A boxplot showed that date hubs were significantly more involved in inter-modular interactions than party hubs (Fig 4a). In robustness tests of the N^0^ and N^+^ networks (Fig 4b and c), the average PCC of inter-modular edges interacting with date hubs was significantly lower than the average PCC of interactions generated in random sampling (658 samples with 100,000 random times). In addition, compared with party hubs, date hubs showed obviously higher degrees and betweenness, such as HSP90AA1, ERBB2, and VEGFA in the top 10 intermodular edge-enriched pathways (S2a Fig), and MAPK3 and HSP90AA1 in top 10 intramodular edge-enriched pathways (S2b Fig).

**Fig 4.**
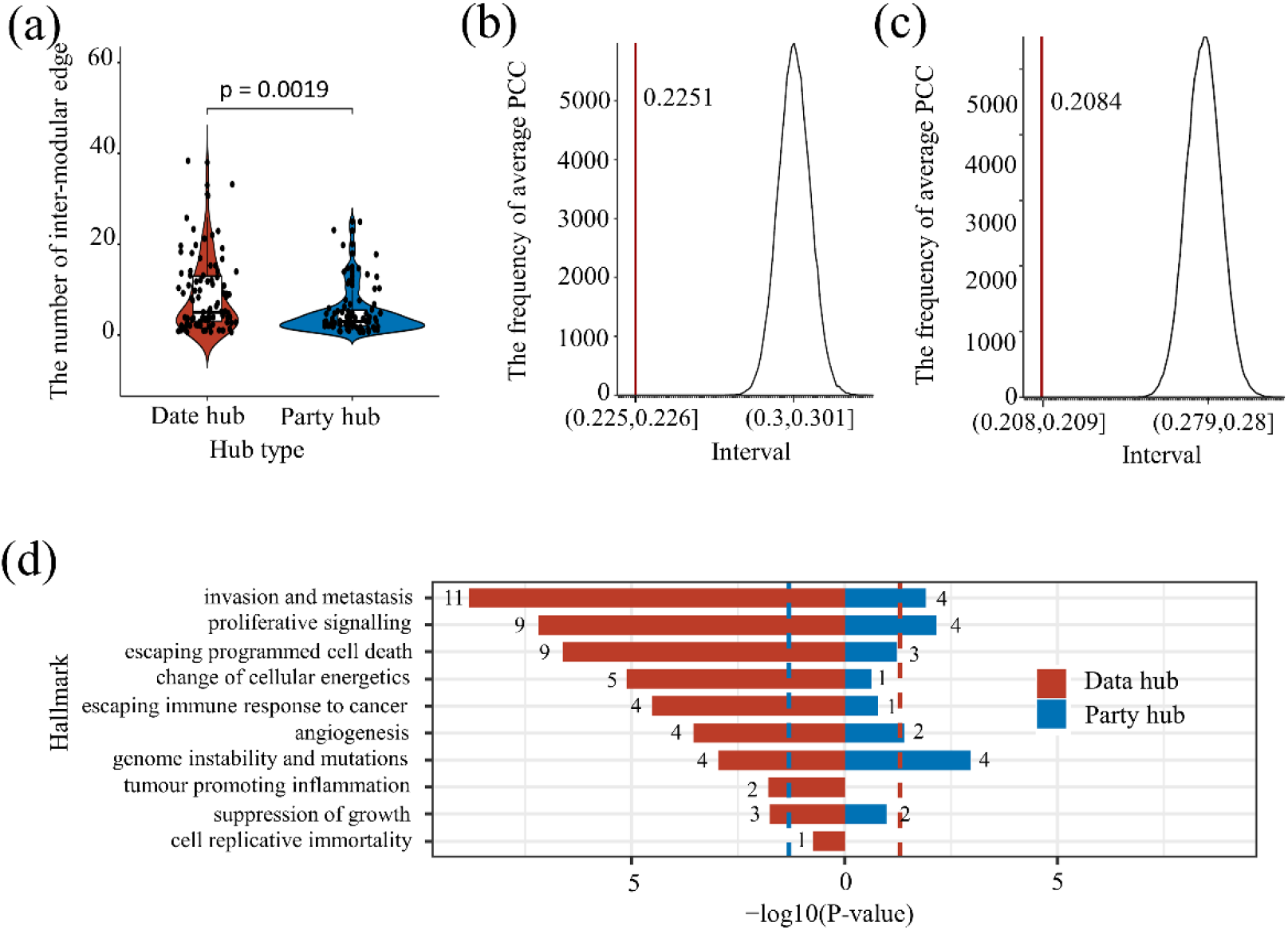
Distribution of date and party hubs and their corresponding KEGG pathways. (a) Statistical comparison of the numbers of intermodular edges consisting of date hubs and party hubs. Sampling tests of intermodular edges in the (b) N^0^ network and (c) N^+^ network. Dashed red lines correspond to the average weight values of inter-modular edges. (d) Cancer hallmark enrichment analysis for date hubs and party hubs. The blue and red dotted lines represent the threshold of -log10 (0.05) calculated by the hypergeometric distribution. The number of genes is marked on the histogram.

Gene expression changes in cancer cells are related to a limited set of special characteristics, often termed cancer hallmarks.[47] We further assessed whether date hubs and party hubs inferred different clinical outcomes. Cancer hallmark enrichment analysis of these hub genes was performed, based on the COSMIC manual annotation of hallmark identification. The hypergeometric distribution was used to infer the significance of the hub genes in cancer hallmarks, with all genes as the background to compute the P*-*value. Overall, date hubs were more highly enriched cancer hallmarks than party hubs, suggesting that date hubs may play a more important role in driving the occurrence and progression of cancer (Fig 4d). Among the 10 cancer hallmarks, only cell-replicative immortality was not enriched for date hubs. The most significant cancer hallmark for date hubs was invasion and metastasis, followed by proliferative signaling and escaping programmed cell death. In contrast, the party hubs showed that “genome instability and mutations” is the most significant cancer hallmark. Specifically, there are nine date hubs enriched in the “invasion and metastasis” hallmark: *GATA3, DDB2, FGFR1, RAC1, ERBB3, ERBB2, GATA2, RET, CDH1, NF2*, and *FGFR2*. However, there are only four party hubs enriched in the “invasion and metastasis” hallmark: *FOXA1, CUX1, PDGFRA*, and *PRKARIA*. The above analysis recapitulated the roles of date hubs and party hubs in terms of hallmark enrichments. This suggests that date hubs are more relevant in the cancer context, especially with regard to invasion and metastasis.

### Core network module of lymph node metastasis in breast cancer

Both date hubs and inter-modular edges describe how protein-protein interaction (PPI) networks change, but date hubs introduce invariance in the dynamic evolution of the network and drive cooperative intermodular interactions. Accordingly, the “core network module” was constructed by connecting date hubs with intermodular edges. We suggest that this core network module contains highly dynamic regions that reorganize to drive or respond to lymph node metastasis in breast cancer (Fig 5a). To evaluate the correlation between this core module and the LNM process, four types of analysis were performed: i) we determined the number of nodes in the core network related to breast cancer progression or metastasis, ii) we calculated the TFC score to detect key interactions, iii) we performed pathway enrichment analysis, and iv) we performed mutation analysis.

**Fig 5.**
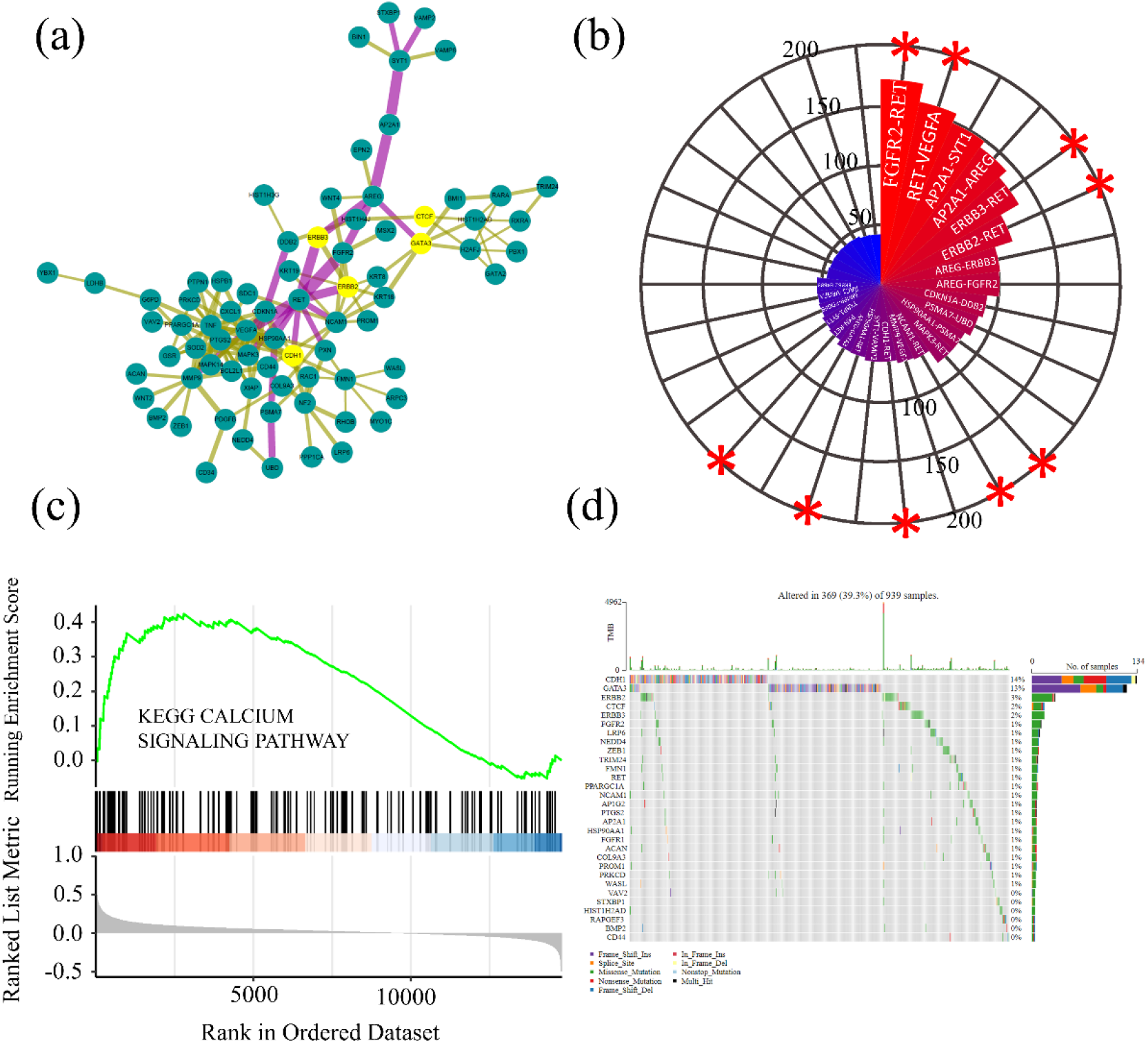
Molecular characteristics of the “ ore network module “. (a) The core network module consists of 76 date hubs and 147 inter-molecular edges. Edges are weighted by TFC scores, and the top 20 edges with the highest TFC scores are denoted in purple. Date hubs with the highest mutation frequency are denoted as yellow nodes. (b) Polar area diagram of network edge prioritization by the TFC scores in the core network module. Asterisks indicate RET-involved interactions. (c) GSEA analysis revealed that the genes of the calcium signaling pathway were enriched across the whole expression profile, and the enrichment status of the calcium signaling pathway was not affected by supervisor selection of differential genes. (d) Mutated date hubs (rows, top 30) are ordered by mutation rate samples (columns) are arranged to emphasize mutual exclusivity among mutations. The row on the right shows the mutation percentage, and the top histogram shows the overall number of mutations. The color coding indicates the mutation type.

For the breast cancer LNM-related core module, most genes (68 of 76) are reported to be associated with breast cancer-related genes, while almost half of the genes (36/76) are related to lymph node metastasis in breast cancer, according to the PubMed database. Additionally, a new score, named TFC, was defined as an edge parameter that was obtained by integrating edge betweenness and the GO semantic similarity of interactions. Here, we ranked the importance of each interaction according to the TFC score. As listed in Table 2, among the top six interactions (FGFR2-RET, RET-VEGFA, AP2A1-SYT1, AP2A1-SYT1, ERBB3-RET, and ERBB2-RET), four are rearranged during transfection (RET)-related, including interactions with two metastatic breast cancer hallmark genes (*ERBB2* and *ERBB3*) (Fig 5b). This interesting finding suggested a key role of RET, as it may be involved in key interactions that need further investigation.

**Table 2.** The 20 interactions with the highest TFC scores in the core network module.

| Proteins | Proteins | TFCs |
| --- | --- | --- |
| FGFR2 | **RET** | 173.0058 |
| **RET** | VEGFA | 157.5393 |
| AP2A1 | SYT1 | 149.0898 |
| AP2A1 | AREG | 143.0134 |
| ERBB3 | **RET** | 134.7705 |
| ERBB2 | **RET** | 117.165 |
| AREG | ERBB3 | 101.4081 |
| AREG | FGFR2 | 100.4558 |
| CDKN1A | DDB2 | 90.53797 |
| PSMA7 | UBD | 87.06091 |
| HSP90AA1 | PSMA7 | 82.14306 |
| MAPK3 | RET | 82.12968 |
| NCAM1 | RET | 69.61411 |
| MMP9 | VEGFA | 68.23644 |
| CDH1 | RET | 65.6743 |
| SYT1 | VAMP2 | 65.67153 |
| HSP90AA1 | RET | 63.11788 |
| AREG | GATA3 | 62.60333 |
| PXN | RET | 58.36475 |
| STXBP1 | SYT1 | 56.96024 |

Furthermore, KEGG pathway enrichment analysis was performed on the core network module. We found that five genes (*VEGFA, FGFR2, RET, ERBB3*, and *ERBB2*) involved in RET-related interactions were enriched in the calcium signaling pathway (P = 0.0109). To verify whether these genes could affect breast LNM through the calcium signaling pathway, we used the GSEA algorithm to evaluate the status of the calcium signaling pathway in the whole expression profile. The results showed that these genes tended to be up-regulated in the calcium signaling pathway, with a P-value of 0.0089 (Fig 5c).

Next, we investigated the genomic mutational signatures, which provided clues for the TNM of breast cancer. The somatic mutations in all samples were compared between date hubs and party hubs. Fig 5c and S3a Fig show the mutation spectrums of highly mutated date-hub and party-hub genes in breast tumor samples. Overall, the mutation frequency of date hubs (39.3%) was higher than that of party hubs (23%). Except for the two most highly mutated genes (*CDH1* and *GATA3*; > 10%), among the date hubs, three more genes (*ERBB2, ERBB3*, and *CTCF*) had relatively high mutation frequencies. In addition, *FGFR2* was a low frequency mutated gene, but the important role of its mutations in breast cancer has been reported.[48]

Accordingly, the above analysis suggested the importance of the sub-network, consisting of RET and its connected genes, as shown in S3b Fig. First, the sub-network contained four of the six most important interactions with highest the TFC scores. Second, the sub-network contained all five calcium signaling pathway-related genes. Third, seven of ten genes in the sub-network had important mutation information, including the core gene of *RET*. By focusing on the sub-network, we further suggested that RET-ERBB2, RET-ERBB2, and RET-FGFR2 interactions are three key interactions for breast cancer LNM.

### Structure-based assessment of the effect of mutations in RET interactions

The occurrence and development of cancer is not only related to changes in expression levels but also to somatic mutations in the coding regions of key genes and their interaction partners. The dynamic assembly of protein complexes is a central mechanism of many cell signaling pathways, which may be regulated by their mutations. To characterize the effect of mutations in the RET interactions, structure models of RET-ERBB2, RET-ERBB3, and RET-FGFR2 complexes were first constructed. The experimental structural data of each protein were collected from the PDB database (PDB id: 6NEC for RET, PDB id: 3PP0 for ERBB2, PDB id: 6OP9 for ERBB2, and PDB id: 2PSQ for FGFR2). Using PRISM, we found that all three interactions can form stable protein complexes, with binding energies of -44.88, -51.97, and -46.33 kcal/mol for the RET-FGFR2, RET-ERBB2, and RET-ERBB3 protein complexes, respectively (Table 3). The structural models for the three protein complexes and their key interfacial residues are shown in S4 Fig. We then mapped missense mutations to the obtained structural models. It can be observed that only the mutation of the D769 in ERBB2 is located at the RET-ERBB2 interface. The distribution of other mutations is relatively scattered over the whole structure.

**Table 3.** Effects of RET mutations on the stability of protein-protein interactions.

| Complex | $\Delta\Delta G_{wt}$ (kcal/mol) | RET Mutation | $\Delta\Delta G_{mt}$ (kcal/mol) | $\Delta\Delta G$ (kcal/mol) |
| --- | --- | --- | --- | --- |
| RET-FGFR2 | -44.88 | L846I | -45.18 | -0.3 |
|  |  | L963V | -44.44 | 0.44 |
|  |  | V778I | -44.25 | 0.63 |
| RET-ERBB2 | -51.97 | L846I | -60.36 | -8.39 |
|  |  | L963V | -55.49 | -3.52 |
|  |  | V778I | -51.97 | 0 |
| RET-ERBB3 | -46.33 | L846I | -55.8 | -9.47 |
|  |  | L963V | -41.3 | 5.03 |
|  |  | V778I | -55.81 | -9.48 |

Footnote: *wt* represents the wild-type protein complex, and *mt* represents the mutant protein complex.

Next, we analyzed the effects of mutations on the stability of protein–protein interactions by calculating the binding affinity. From the TCGA sample, we found three mutations in RET: the tyrosine domain V778I mutation; and two kinase domain mutations, L846I and L963V. As shown in Table 3, the change in the binding energy (ΔΔG) of the three mutations (V778I, L846I, and L963V) on the RET-ERBB2, RET-ERBB3, and RET-FGFR2 complexes shows that their binding energy may become more stable or reduce slightly. Similar results were also found for mutations of ERBB2, ERBB3, and FGFR2 (S1 Table), and even for the D769H(Y) mutation located at the RET-ERBB2 interface. In conclusion, the structural modeling, *in silico* mutagenesis, and comparison of the predicted binding energies revealed that the mutations did not disturb protein-protein interactions, suggesting other kinds of mechanisms.

In addition, we used a structure-based statistical mechanical model implemented in AlloSigMA[45] to obtain a direct estimate of the allosteric effects across protein-protein interactions caused by these missense mutations. As shown in S5-S8 Fig, except for V778I in RET, almost all other mutations have no allosteric effects on their protein partners. We observed that V778I in RET showed allosteric effects on all its partners, including ERBB2, ERBB3, and FGFR2 (Fig 6a). The V778I mutation is a large amino acid, while in AlloSigMA, it is defined as an UP mutation. The allosteric energy profile showed that the energy of residues near V778 was negative and relatively low, which demonstrated the greater local stability of this region. However, for the interaction partners of RET, the allosteric free energy of residues was generally positive and relatively high, indicating that the interaction partners became unstable above the V778I mutation in RET. In detail, we predicted that FGFR2 had an obvious peak (0.99kcal/mol) at residue P582 (Fig 6b), and ERBB2 had a local increase in kinetics at residues E719, I740 to V746, D873, E874, H878, I886, L891 to L895, I926, E930 to R940, and I961(Fig 6c). In addition, we observed a distal effect on the ERBB3 activation loop (P842-K853) and residues G713 to S715, where the contact network became unstable (Fig 6d). Overall, our results showed that the V778I mutation in the RET protein had an unstable effect on the contact network in its interacting partners. This may affect the interaction through allosteric communication between RET and its interacting partners.

**Fig 6:**
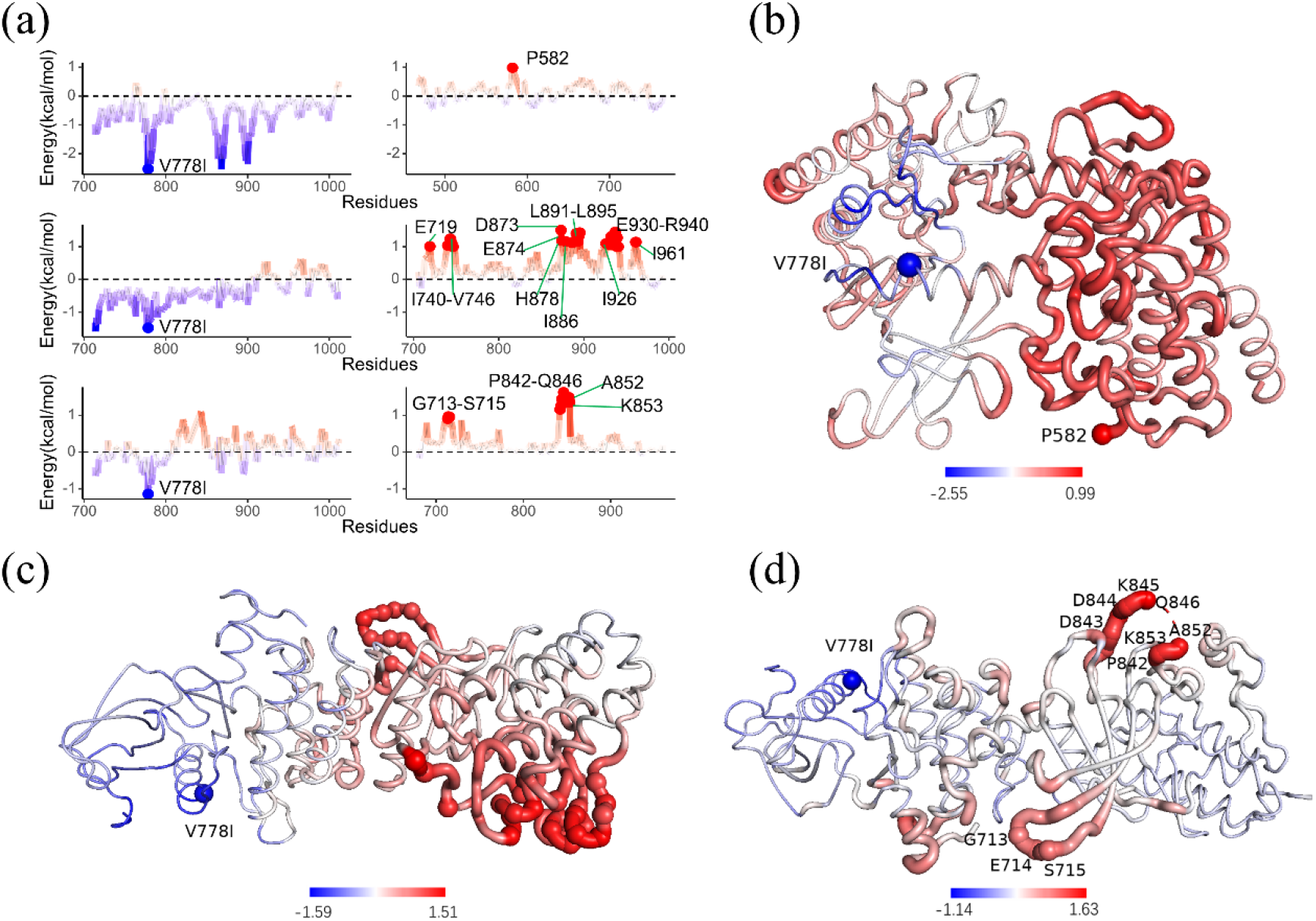
Allosteric effects caused by V778I in RET. (a) Allosteric energy profiles for V778I in RET in three protein complexes, predicted by AlloSigMA. Mapping of allosteric energy on 3D structures of the (b) RET-FGFR2, (c) RET-ERBB2, and (d) RET-ERBB3 interactions.

## Discussion and Conclusion

Cancer metastasis is therefore an evolving disease and the combined outcome of cells that metastasize and a series of microenvironmental factors that they interact with, collude, or surmount. Although each instance of metastasis could be unique, the quest is to find commonalities that could be targeted therapeutically. The complexity of metastasis through its chronological progression, and its manifestation in various biological scales, calls for a systems approach to understand metastasis mechanistically. In this work, we proposed a method that combines differential modular analysis with mutational structural analysis to study the dynamic process and molecular mechanism of cancer metastasis. By applying this method to study the LNM of breast cancer, we identified some key factors, including the core network module, key PPIs, and a potential allosteric mutation.

From the methodology, we combined gene co-expression data, PPI networks and structures, and genetic variation together to understand disease progression at both the systems and molecular levels. First, we constructed two PPI networks with the same topology but weighted with different gene co-expression data. Then, different modular structures were obtained by modular analysis and the two PPI networks associated with different disease states were compared. The third and fourth steps represent the main idea of our method, that inter-modular edges and date hubs obtained from different modular analysis afforded more functional contexts for disease progression. The network module associated with disease progression was constructed by connecting date hubs with inter-modular edges. Based on the core network module, subsequent analysis included ranking all the interactions by TFC scores and mutation analysis. Lastly, the structures of PPIs were modeled and mutations with important functions were predicted.

In summary, the novelty of our method was two-fold. First, differential network analysis methods were normally node- and edge-based, or they involved a comparison of the global topology. Our method here was module-based, which could help in discovering shared or changed functional patterns. The differential modular analysis lies somewhere between node/edge analysis and global topological differential network analysis, shedding more light on the underlying mechanisms of biological systems. Second, the proposed method was novel insofar as it linked allosteric mutation with PPI network analysis to characterize the differentially interacting modules. As such, it could identify key PPIs and active mutations during LNM. This idea has been used previously to study the role of Bcl-2 proteins in breast cancer.[49]

In breast cancer, the activation mutation in the ERBB2 is a well-known oncogenic driver. The interaction of ERBB2 with other protein partners (or dimerization) activates various oncogenic signaling pathways related to breast cancer metastasis, including the Smad2/3, RAS/RAF/MAPK, PKC, and PI3K-Akt signaling pathways.[50] Recent computational structural modeling with biochemical and cell biological analyses suggests that ERBB2 mutations need to mutate ERBB3, and then promote oncogenesis and invasion of breast cancer via PI3K pathway activation.[51]

However, we identified that calcium signaling pathways could be considered an ERBB2 participation noncanonical pathway related to LNM in breast cancer. As a second messenger, the intracellular calcium ion (Ca^2+^) plays direct and robust roles in many biological processes. Several studies have reported that calcium signaling pathways are essential to cancer progression. In particular, calcium signaling pathways regulate key processes, from inflammation to apoptosis, that are involved in breast cancer tumorigenesis,[52] metastasis,[53] and resistance to chemotherapy.

The current standard of care for the treatment of metastatic ERBB2 breast cancer is the combination of seven inhibitors: trastuzumab, pertuzumab, trastuzumab emtansine, trastuzumab deruxtecan, lapatinib, neratinib, and tucatinib.[54] However, drug resistance is common and remains a major unresolved clinical problem. Targeting the interaction between ERBB2 and a number of non-canonical RTKs (i.e. EGFR, ERBB3, and ERBB4) has emerged as a promising therapeutic method that overcomes drug resistance in treating breast cancer metastasis.[55]

Here, we considered that breast cancer LNM may be the outcome of allosteric driver mutations in RET.[56] As one of the RTK members, RET is a target for several kinds of human cancer, such as thyroid, breast, and colorectal carcinoma. Many RET missense mutations are known to be causally associated with breast cancer, including extracellular domain mutations C611R, C620F, L633V, C634R, C634F, and T636M, and the kinase domain M918T.[57] A structural understanding of the mutational roles of RET in different interactions suggested V778I as an allosteric mutation, raveling potential targets to prevent LNM in breast cancer from overcoming resistance in ERBB2.

Altogether, the analysis characterized a differential module representing significant changes in their interaction patterns during LNM in breast cancer, and their functions roles in terms of three enrichment analyses and mutation analysis based on the structural level. In addition, the differential module is potentially valuable not only for understanding LNM but also for developing effective diagnosis, prognosis, and treatment strategies.

## Supporting Information

S1 Table. The effects of FGFR2, ERBB2 and ERBB3 mutations on the stability of protein-protein interaction.

S1 Fig. (a) Volcano map of the lymph node metastasis group and the non-lymph node metastasis group in breast cancer. (b) Kaplan-Meier survival curves of the lymph node metastasis group and the non-lymph node metastasis group. (c) The heatmap of Jaccard similarity coefficients for the modules between the N^0^ and N^+^ PPI networks. (d) The modular structure of N^+^ network.

S2 Fig. The distribution of degree and betweenness for hubs involved in inter-modular edge and intra-modular edge enriched top 10 Reactome pathways.

S3 Fig. The oncoplot for top 30 mutated party hubs and *RET*-centered sub-module.

S4 Fig. The structural models for the RET-FGFR2, RET-ERBB2, and RET-ERBB3 protein complexes and the distribution of missense mutations.

S5 Fig. The allosteric effect of RET mutations on its three interaction partners. S6 Fig. The allosteric effect of FGFR2 mutations on RET.

S7 Fig. The allosteric effect of ERBB2 mutations on RET. S8 Fig. The allosteric effect of ERBB3 mutations on RET.

## Acknowledgments

This work was supported by the National Natural Science Foundation of China (31872723), a Project Funded by the Priority Academic Program Development (PAPD) of Jiangsu Higher Education Institutions, the China Postdoctoral Science Foundation (2016M590495), and the Jiangsu Planned Projects for Postdoctoral Research Funds (1601168C).

## Author Contributions

The manuscript was written through contributions of all authors. All authors have given approval to the final version of the manuscript. ^*¶*^These authors contributed equally.

## Notes

The authors declare no competing financial interest.

## Abbreviations

LNM: lymph node metastasis
RET: rearranged during transfection
PPIN: protein-protein interaction network
EGFR: Epidermal growth factor receptor
BRCA: breast cancer
DEGs: differentially expressed genes
aPCC: average Pearson’s correlation coefficient
CPL: characteristic path length
TFC: topological-functional connection
TCGA: The Cancer Genome Atlas
KEGG: Kyoto Encyclopedia of Genes and Genomes
GSEA: Gene set enrichment analysis
COSMIC: Catalogue of Somatic Mutations in Cancer
PDB: Protein Data Bank
PRISM: Protein Interactions by Structural Matching.

